# CCM2 deficient endothelial cells undergo a mechano-dependent reprogramming into senescence associated secretory phenotype used to recruit endothelial and immune cells

**DOI:** 10.1101/2021.02.22.432204

**Authors:** Daphné Raphaëlle Vannier, Apeksha Shapeti, Florent Chuffart, Emmanuelle Planus, Sandra Manet, Paul Rivier, Olivier Destaing, Corinne Albiges-Rizo, Hans Van Oosterwyck, Eva Faurobert

**Author notes:** Deceased (Sandra Manet). Co-last authors (Corinne Albiges-Rizo, Hans Van Oosterwyck, Eva Faurobert).

## Abstract

Cerebral Cavernous Malformations (CCM) is a cerebrovascular disease in which stacks of dilated haemorrhagic capillaries form focally in the brain. Whether and how defective mechanotransduction, cellular mosaicism and inflammation interplay to sustain the progression of CCM diseases is unknown. Here, we reveal that CCM1- and CCM2-silenced endothelial cells enter into senescence associated with secretory phenotype (SASP) that they use to invade the extracellular matrix and attract surrounding wild-type endothelial and immune cells. Further, we demonstrate that this SASP is driven by the mechanical and molecular disorders provoked by ROCKs dysfunctions. By this, we identify CCM1/2 and ROCKs as parts of a scaffold controlling senescence, bringing new insights into the emerging field of the control of aging by cellular mechanics. This discovery reconciles the dysregulated traits of CCM1/2-deficient endothelial cells into a unique mechano-dependent endothelial fate that links perturbed mechanics to microenvironment remodelling and long-range activation of endothelial and immune cells.

## Introduction

Cerebral Cavernous malformations (CCM) are stacks of overgrown, dilated and haemorrhagic venous capillaries formed by a unique layer of poorly joined endothelial cells (EC) without intervening cerebral mural cells(1). Loss-of-function mutations on 3 genes (*CCM1/KRIT, CCM2/*Malcavernin, *CCM3/PDCD10*) are associated with the familial form of the disease(2,3). *CCM1* or *CCM2* associated disease develops later in life than CCM3 which is a more aggressive form of the disease(4,5).

CCM lesions expand with time and they become infiltrated by immune cells that sustain a chronic inflammatory response(6,7). Intriguingly, CCM lesions are composed of a mosaic of mutant and wild-type EC (8–11). Malinverno and colleagues have further shown that the majority of EC bordering large mature ccm3 caverns are actually wild-type EC that have been attracted to the lesion site at least in part by mutant EC(11). Other studies have reported that *CCM* mutant EC secrete metalloproteases(12–14) or cytokines(15), over-produce ROS(16), present defective autophagy(17) or that they undergo an endothelial to mesenchymal transition (EndMT)(18). Moreover, loss of CCM proteins activates β1 integrin (19,20), p38 MAPK(21), ERK5-KLF2/4(20,22,23) and TLR4 signaling pathways(24). However, how these various dysregulations interplay to generate CCM lesions is not well known.

A remarkable feature of CCM lesions is their peculiar mechanical microenvironment. Indeed, EC in CCM lesions experience disturbed forces coming from stagnant blood flow(25) on their luminal side and increased ECM stiffness upon matrix remodelling on their basal side(19). Increased RhoA/ROCK-dependent intracellular tension (19) is a conserved feature of *CCM* mutant EC in humans and animal models (13,26–28). ROCK over activation stimulates the polymerization of a contractile acto-myosin cytoskeleton that shifts the tensional homeostasis between cell-cell and cell-extracellular matrix (ECM) adhesions. We previously showed that the endothelial tensional homeostasis is actually under the control of the coupled activities of the two ROCK isoforms(29). The molecular scaffold formed by the association of CCM1 and CCM2 recruits ROCK2 to VE cadherin-complexes to promote the polymerization of a cortical acto-myosin cytoskeleton supporting cell-cell junctions. At the same time, this CCM/ROCK2 complex keeps ROCK1 kinase activity low thereby limiting the adhesion of the cell to the ECM. When the CCM1/2 complex is lost, ROCK2 delocalizes from VE-cadherin while ROCK1 gets over activated and promotes the polymerization of numerous β1 integrin-anchored acto-myosin stress fibers that most likely tear the cell-cell junctions apart(29). Importantly, it is yet unknown whether, beyond their role on the architecture of the endothelium, ROCKs are also involved in the control of gene expression downstream of CCM2.

The mechanical defects play a primary role in the development of the disease. Inhibition of both ROCKs with chemical inhibitors, among which fasudil, blocks the genesis and maturation of CCM lesions in animal models(30–32). However, the toxicity of these drugs precludes their use in patients. It is therefore critical to find new therapies targeting specific downstream pathways. Toward this goal, we need to find a mechanistic explanation that could integrate all the different dysregulated traits of *CCM* mutant EC.

Here, we reveal that the transcriptome of CCM2-silenced EC presents a signature of Senescence Associated Secretory Phenotype (SASP). Cellular senescence contributes to a wide variety of human age-related pathologies, including cancer, fibrosis, cardiovascular diseases, or neurological disorders(33). Further, we demonstrate that CCM2-silenced EC indeed enter into premature senescence and acquire degradative and invasive skills that stimulate angiogenesis *in vitro*. CCM2-deficient EC gain paracrine functions through secreted factors that attract wild-type EC and immune cells. Remarkably, we show that this SASP is a mechano-dependent process triggered by dysfunctional ROCK1 and ROCK2 and by increased EC contractility. By this, we identify CCM1/2 proteins and ROCKs as part of a mechanotransduction scaffold controlling senescence. This unexpected endothelial fate transition triggered by the loss of CCM2 unifies all the known dysregulated features of CCM2-deficient EC and establishes a new molecular mechanism supporting the mosaicism of the CCM lesions and their inflammatory state.

## Results

### 1) The loss of CCM2 turns on a SASP transcriptomic program in endothelial cells

KRIT and CCM2 proteins interact with each other to form a molecular scaffold(34). We previously showed that, owing to the stabilizing effect of CCM2 on KRIT, both proteins are lost when CCM2 is silenced(19). We therefore chose to silence CCM2 in order to deplete the entire KRIT/CCM2 complex. To study the gene expression program of a pure population of EC depleted for CCM2, we performed RNA sequencing on monolayers of human umbilical vein endothelial cells (HUVEC) after two consecutive rounds of transfection with CCM2 targeting siRNA (siCCM2) or with non-targeting siRNA (siNT). CCM2 silencing was of 86% (figS1A) as reported in Lisowska et al., 2018(29). A total number of 2057 differentially expressed genes (DEGs) (fold change [FC] ≥ 2; *P* < 0.05, FDR corrected using Benjamini Horchberg method (35)) were identified in siCCM2-treated compared to siNT-treated HUVEC among which 1318 genes were upregulated while 739 were downregulated (Table S1A).

To investigate the cellular functions altered by the loss of CCM2, we performed a Gene Ontology analysis on these DEGs using cellular component (Fig 1A) and biological process (Fig 1B) annotations. Up-regulated genes were associated with the plasma membrane, secretory vesicles, extracellular matrix and focal adhesions (Fig 1A). They related to ECM organization, cell adhesion and migration, secretion, inflammatory response to cytokines and calcium homeostasis (Fig1B). Down-regulated genes were associated with the nuclear part of the cell, chromosomes, chromatin, the mitotic spindle pole and kinetochores and microtubules (Fig 1A). They relate to DNA replication, recombination and repair, chromosome segregation, microtubule-dependent movements and cell cycle progression (Fig 1B). A Reactome analysis further confirmed these results (Fig 1C). In fact, these up and down-regulated functions are characteristic of a striking unique cellular state; a senescence-associated with secretory phenotype (SASP) (Fig 1C). This phenotype defines the ability of cell-cycle-arrested cells to secrete pro-inflammatory cytokines, chemokines, growth factors and proteases giving rise to ECM remodelling and to the stimulation of neighbour cells proliferation and invasiveness(36).

**Figure 1.**
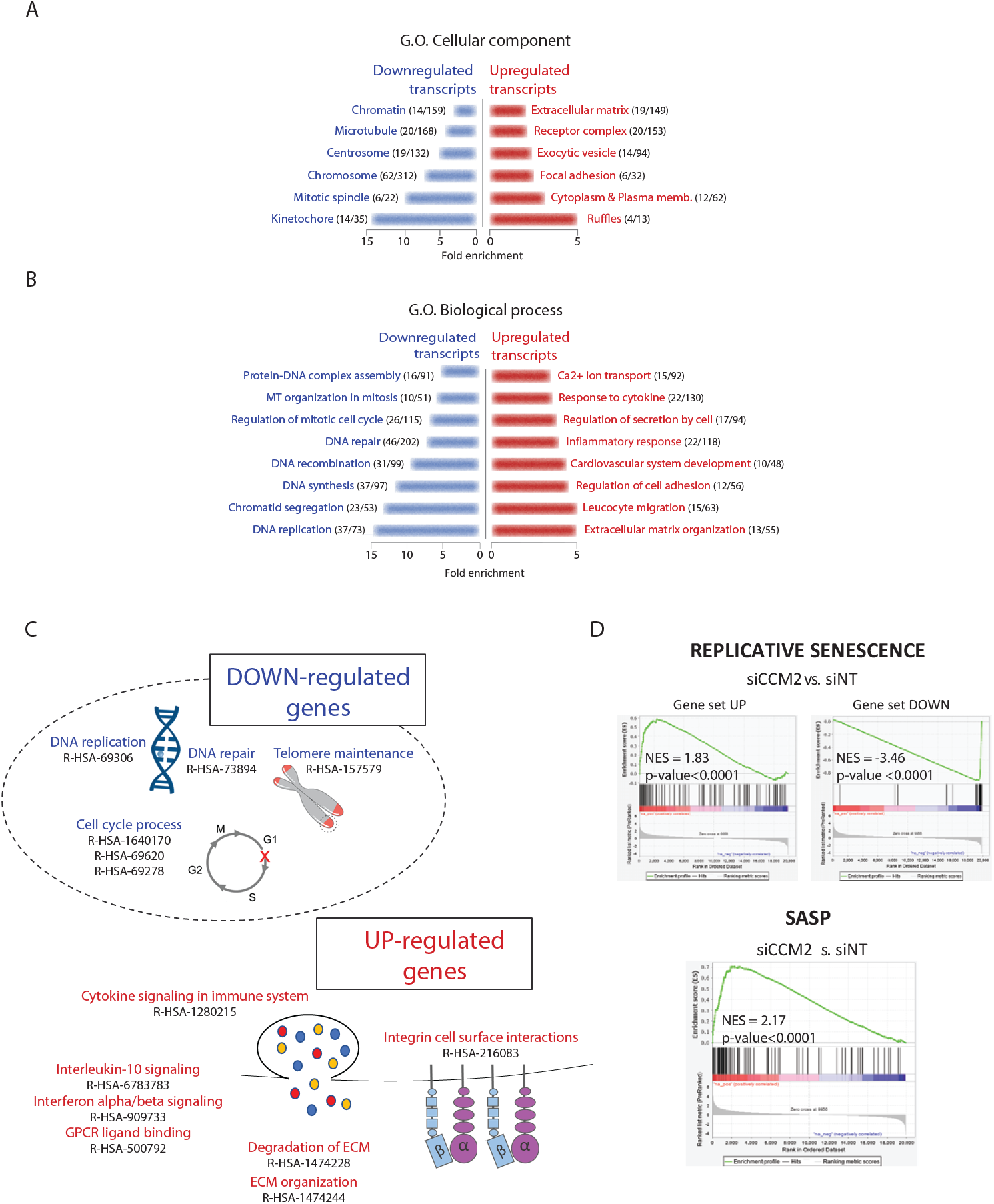
The loss of CCM2 turns on a SASP transcriptomic program. (A)Gene Ontology enrichment analysis of cellular components in downregulated and upregulated genes in siCCM2 HUVEC compared to siNT HUVEC, bar graphs represent the fold enrichment. (B) Gene Ontology enrichment analysis of biological functions in downregulated and upregulated genes in siCCM2 HUVEC compared to siNT HUVEC, bar graphs represent the fold enrichment. (C) Schematic representation of the Reactome analysisof enriched pathways in siCCM2 HUVEC. (D) GSEA profiles showing a significant normalized enrichment score (NES) of gene sets associated with replicative senescence(37) and SASP(39) in siCCM2 HUVEC transcriptome.

To comfort the hypothesis that a SASP transcriptomic program is turned on in siCCM2 HUVEC, we search for specific transcriptomic signatures of senescence and SASP in the literature corresponding to gene sets enriched in senescent cells among which fibroblasts and endothelial cells(37–40). These gene sets relate to up- and down-regulated genes (Table S2). We then searched for an enrichment in these gene sets in the CCM2-depleted transcriptome using Gene Set Enrichment Analysis (GSEA). We confirmed that these premature replicative senescence and SASP signatures are significantly enriched in the CCM2-silenced transcriptome (Fig1D, Fig S2). Overall, these functional analyses of the transcriptome suggest that EC undergo a SASP when CCM2 is lost.

### 2) CCM2- and KRIT-depleted EC display hallmarks of SASP

Since a SASP transcriptomic program is turned on upon the loss of CCM2, we next sought for features of premature senescence in these cells. We looked for different hallmarks, as the combination of multiple traits is required to ascertain senescence(41). HUVEC were analysed at passage 4 when siNT cells are still proliferative and healthy. In addition to flattening and elongating upon the production of transversal stress fibers (Fig S3A), CCM2-depleted EC expressed almost a 3-fold increase in lysosomal senescence-associated β-galactosidase (SA-β-gal) activity, the historical marker of senescence (Fig 2A). In addition, their nuclei displayed senescence-associated heterochromatin foci (SAHF) as revealed by spots in DAPI and HIRA staining (Fig 2B) and their area was increased (Fig 2C). As senescence leads to cell cycle arrest, we looked at the expression level of the cell cycle inhibitors CDKNs. Among them, p21/CIP1 and p15/INK4b were 3-fold upregulated. On the contrary, cyclin dependent kinase 1, its regulator CKS1 and cyclin 2A as well as the transcription factor E2F1, a driver of S phase entry, were all dramatically downregulated (fig S1B). Using BrdU incorporation, we detected a 2-fold reduction in the percentage of cells in S phase and an accumulation in G1 indicative of a defect in the G1/S transition of the cell cycle (Fig 2E). This translated into a significantly lowered rate of proliferation of the EC population as shown by impedance measurements (Fig 2F), and a two-fold lowered percentage of cells positive for the proliferative marker Ki67 (Fig 2G). Overall, the combination of all these traits confirms that the loss of CCM2 indeed induced premature senescence in HUVEC.

**Figure 2.**
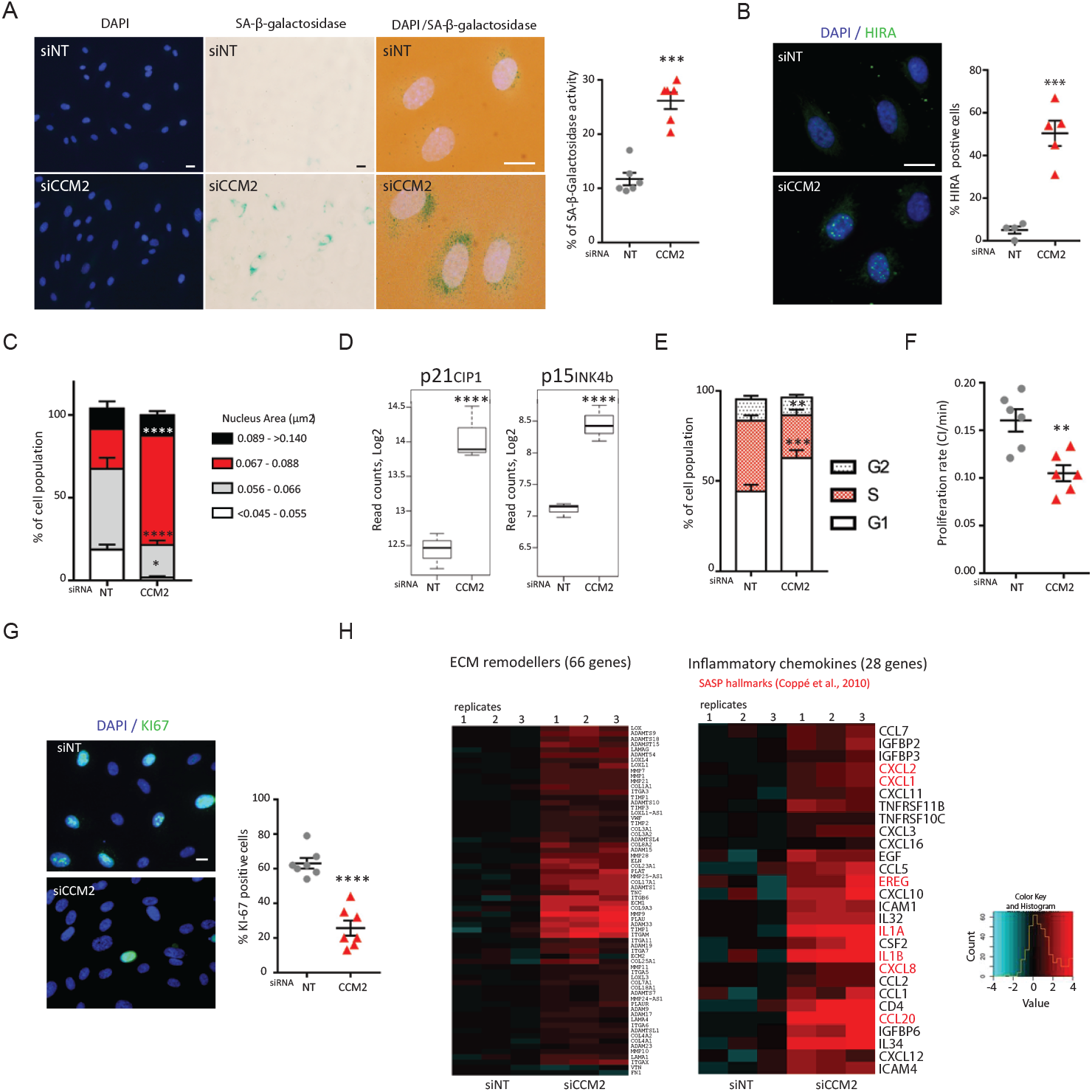
CCM2-depleted EC display hallmarks of SASP. (A)(left) Representative images of DAPI and SA-βgalactosidase staining of siNT and siCCM2 HUVEC. (Middle) Merge of DAPI and SA-β-galactosidase stainings at higher magnification. (Right) Quantification of the % of positive cells for SA-β-galactosidase. Error bars are means ± SEM from 6 independent experiments. (B)(left) Representative images of merged DAPI and HIRA stainings of siNT and siCCM2 HUVEC. (Right) Quantification of the % of HIRA positive cells. Error bars are means ± SEM from 5 independent experiments. (C) Histogram of the cell population in function of their nucleus area. Error bars are means ± SEM from 4 independent experiments. (D) Boxplots of the read counts for p21/CIP and p15/INK4b mRNA. Error bars are means ± SEM from 3 independent experiments. (E) Quantification by BrdU assay of the percentage of cells in each phase of the cell cycle. Error bars are means ± SEM from 8 independent experiments. (F) Proliferation rate of siRNA transfected HUVEC measured by impedance using XCELLigence. Error bars are means ± SEM from 4 independent experiments. (G)(left) Representative images of the proliferation marker Ki-67 staining (green) merged with DAPI staining. (Right) Quantification of the percentage of cells positive for Ki-67 staining. Error bars are means ± SEM from 7 independent experiments. (H) Heatmap of expression of ECM remodelling proteins (left) and of SASP factors (right) over the 2 siRNA conditions, 3 replicates per condition. (*) P-value<0.05; (**) P-value<0.005; (***) P-value<0.0005; (****) P-value<0.00005. Scale bars are equal to 10 µm.

Senescent cells secrete paracrine factors that can promote tumor development in vivo by engaging deleterious inflammatory responses and malignant phenotypes such as proliferation and invasiveness (42). We found that siCCM2 treated HUVEC overexpressed ECM remodelling mediators including matrix proteins, metalloproteases of the MMP and ADAM families, the plasminogen activator uPA and cross-linking enzymes (Fig 2H left). Moreover, they overexpress cytokines and inflammatory chemokines among which IL-1A and B, IL8, CXCL1, 2, CCL20 and EREG that are hallmarks of SASP (43) (Fig 2H right).

Having shown that the loss of CCM2 leads to SASP, we wondered whether this would be a common feature with the loss of KRIT and CCM3. KRIT-depleted HUVEC displayed the same senescent phenotype as CCM2-depleted HUVEC as shown by a significant increase in cells expressing SA-βgal activity (Fig 3A) and SAHF (Fig 3B), a significant decrease in Ki67-positive cells (Fig 3C) and with a lowered rate of proliferation (Fig 3D). Interestingly, CCM3-depleted EC did not display marks of senescence in good agreement with its distinct role in EC biology and onset of the disease(44). Indeed, CCM3-depleted HUVEC behaved as control cells in these assays. They did not show increased SA-βgal activity (Fig 3A), nor SAHF (Fig 3B). They had a normal level of Ki67 positive cells (Fig 3C) and they did not proliferate differently from control cells (Fig 3D). Therefore, consistently with their strong association and regulation of common signaling pathways(19,44), KRIT and CCM2 loss similarly lead to premature cellular senescence.

**Figure 3.**
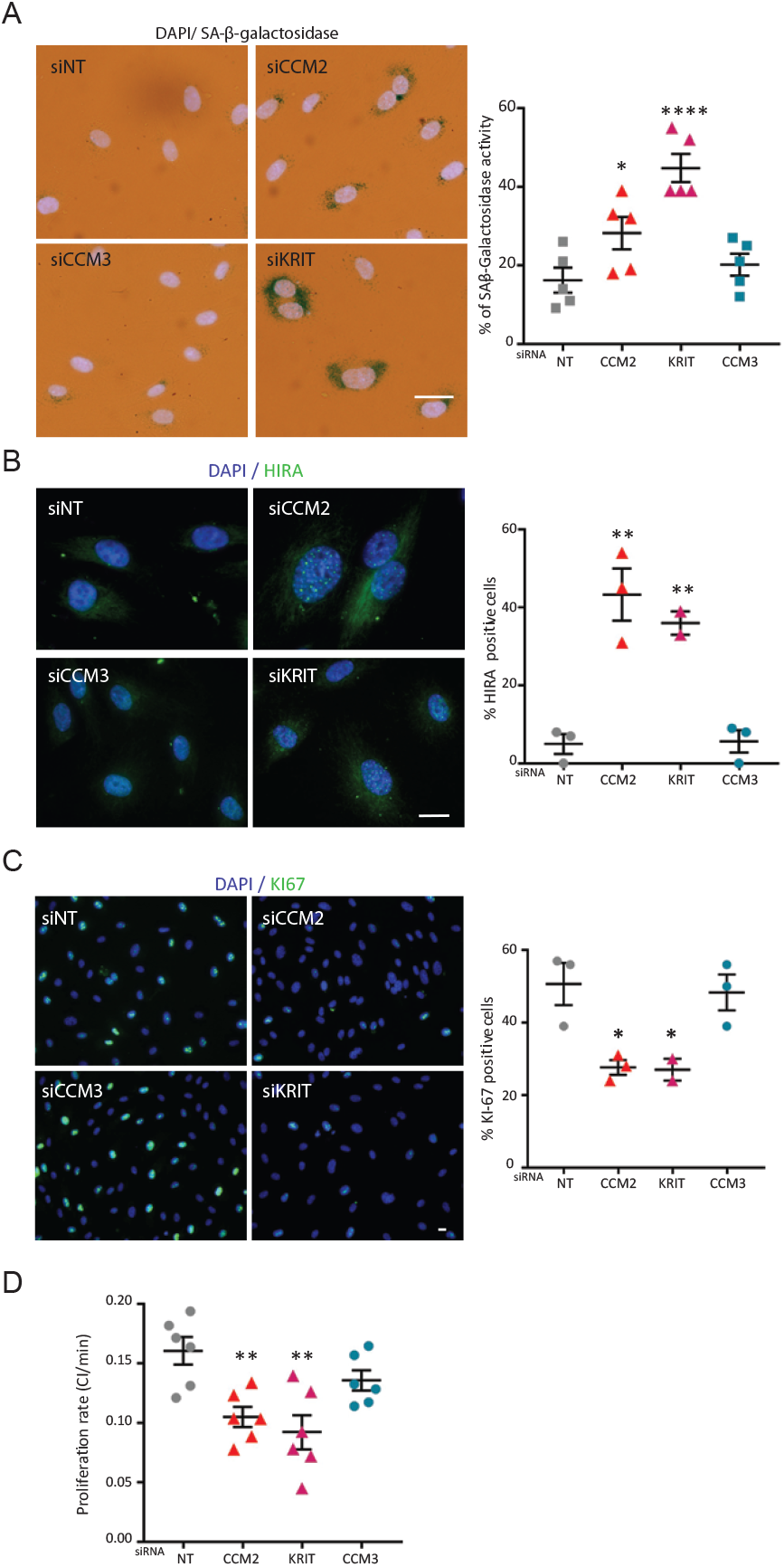
The loss of KRIT similarly to that of CCM2 leads to senescence in EC whereas the loss of CCM3 does not. (A)(left) Representative merged images of DAPI and SA-β-galactosidase stainings. (Right) Quantification of the % of positive cells for SA-β-galactosidase. Error bars are means ± SEM from 5 independent experiments. (B)Representative images of merged DAPI and HIRA stainings. (Right) Quantification of the % of HIRA positive cells. Error bars are means ± SEM from 3 (2 for KRIT) independent experiments. (C) Representative images of the proliferation marker Ki-67 staining (green) merged with DAPI staining. (Right) Quantification of the percentage of cells positive for Ki-67 staining. Error bars are means ± SEM from 3 (2 for KRIT) independent experiments. (D)Proliferation rate of siRNA transfected HUVEC measured by impedance using XCELLigence. Errors bars are means± SEM from 3 independent experiments.

### 3) ROCK2 controls the SASP transcriptomic program of CCM2-depleted EC

ROCK-dependent perturbations in the mechanotransduction of EC have a major role in the genesis and progression of CCM lesion. Inhibition of ROCK is sufficient to block the formation and the maturation of CCM lesions (30–32). We previously showed that the CCM1/2 complex is a scaffold recruiting ROCK2 at VE-cadherin complexes thereby limiting ROCK1 kinase activity to maintain the tensional homeostasis between cell-cell and cell-ECM adhesions and to preserve the integrity of the endothelial monolayer(29). However, it is yet unknown whether ROCKs are also involved in the control of gene expression downstream of CCM2 and in particular in the regulation of this SASP transcriptomic program. Hence, we analysed the contribution of ROCK1 and ROCK2 by performing RNA sequencing on monolayers of CCM2-silenced HUVEC that were additionally silenced for ROCK1 or ROCK2 (Fig S1A) in the same set of experiments as that shown in figure 1. We have previously shown that the additional depletion of ROCK1 but not ROCK2 restores the morphological defects of the CCM2-deficient EC and their permeability barrier(29). Depletion of ROCK1 or ROCK2 alone were performed as controls. Strikingly, 40% of the DEGs with FC>2 (54% of all DEGs) had their expression significantly returned toward the control level by the additional silencing of ROCKs (Fig 4A, Table S1B). The silencing of ROCK2 had a stronger restoring effect than that of ROCK1 on the number of restored genes (Fig 4A) and their level of expression (Fig 4B, Fig S2A). Importantly, silencing of ROCK2 alone had overall an opposite effect to that of CCM2 on gene expression (Fig 4B), suggesting that ROCK2 acquires a gain of transcriptional function when CCM2 is lost.

**Figure 4.**
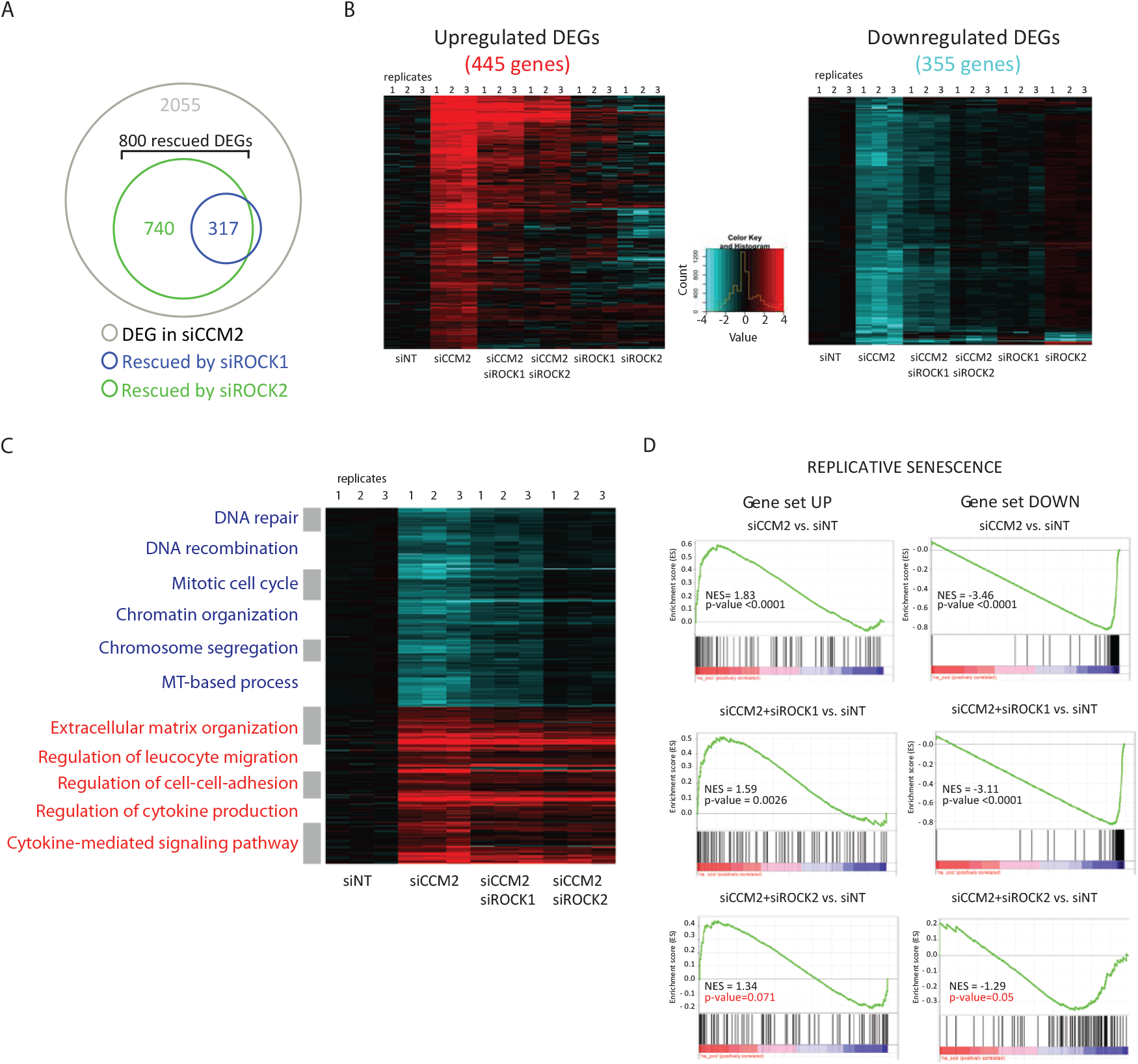
ROCK2 controls the SASP transcriptomic program of CCM2-depleted EC. (A)Venn diagrams showing overlap of DEGs with FC≥ 2; P < 0.05 in siCCM2 HUVEC (grey) with DEG rescued in siCCM2+siROCK1(Blue) siCCM2+siROCK2 (Green). (B) Heatmap of DEGs with FC≥ 2; P < 0.05 in siCCM2 HUVEC rescued by ROCKs. Upregulated (left) and downregulated (right) genes over the 6 siRNA conditions, 3 replicates per condition. (C) Clustered heatmap of GO enriched in up-(left) and down-(right) regulated genes in siCCM2 HUVEC. (D) GSEA enrichment plot showing the lossof significant enrichment in replicative senescence signature in siCCM2+ROCK2 but not in siCCM2+ROCK1 HUVEC transcriptome.

We then studied the restoring effect of ROCK1 or ROCK2 silencing on the biological functions perturbed by CCM2 depletion. Figure 4C shows a clustered expression heatmap of DEGs belonging to the Gene Ontology (G.O) biological processes presented in figure 1B. The expression of DEGs related to the down-regulated nuclear functions was fully restored by ROCK2 depletion while ROCK1 depletion had only a partial effect. In addition, up-regulated peri-membrane functions were rescued by the silencing of ROCK2 to a higher extent than that of ROCK1 (Fig 4C). Going further, we focused on the effect of ROCKs on the signatures of senescence or SASP found in siCCM2 transcriptome. While these signatures of senescence were still present upon ROCK1 depletion, they were not anymore significantly enriched in HUVEC doubly silenced for CCM2 and ROCK2 (Fig 4D, Fig S2B) highlighting the crucial role of dysregulated ROCK2 in the onset of the SASP transcriptomic program.

Overall, our transcriptomic data reveal that, beyond their role on the tensional homeostasis of the endothelial monolayer, ROCK2 and to a lesser extent ROCK1 control the expression of an important fraction of the genes regulated by CCM2. These genes are involved in a transcriptomic program supporting the onset of SASP when CCM2 is lost.

### 4) ROCKs dysfunctions induce premature senescence in CCM2-depleted EC

Having shown that ROCKs control the expression of genes involved in SASP, we next asked whether dysfunctional ROCKs played a causal role in the entry into SASP of CCM2-depleted EC. Additional silencing of ROCK1 or ROCK2 was similarly efficient in preventing the appearance of most of the features tested i.e. SA-β-gal activity, HIRA-positive SAHF and restored normal level of Ki67+ cells (Fig 5A, B, E). However, ROCK2 silencing was more efficient in preventing the accumulation in G1 (Fig 5D), consistently with its higher efficiency in lowering the expression of p21/CIP1 and p15/INK4b (Fig 5C) and in restoring the expression of down-regulated cyclins and cyclin dependent kinases (Fig S1B). Moreover, the silencing of ROCK2 and to a lesser extent that of ROCK1 lowered the expression of SASP factors, i.e. ECM remodelers or inflammatory chemokines (Fig 5F).

**Figure 5.**
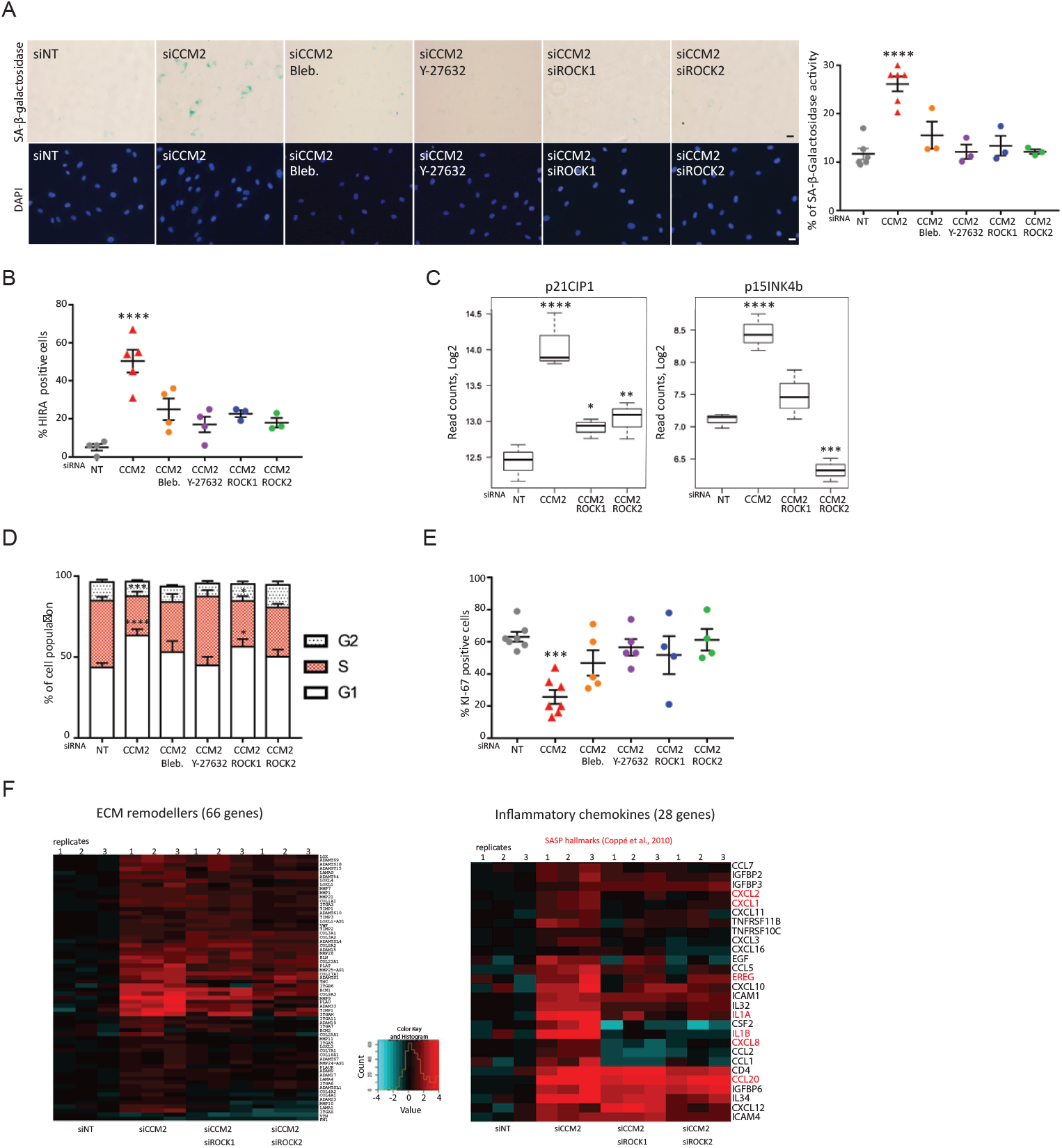
ROCKs dysfunctions induce premature senescence in CCM2-depleted EC. (A)(left) Representative images of SA-β-galactosidaseand DAPI stainings of siRNA-transfected HUVEC or treated with blebbistatin or Y27632 at 10 μM. (Right) Quantification of the % of positive cells for SA-β-galactosidase. Error bars are means± SEM from 3 independent experiments. (B) Quantification of the % of HIRA positive cells in siRNA-transfected HUVEC or treated with blebbistatin or Y27632. Error bars are means ± SEM from 4 (drug treatments) and 3 (ROCKs silencing) independent experiments. (C) Boxplots of the read counts for p21/CIP and p15/INK4b mRNA after depletion of ROCK1 or ROCK2. Error bars are means ± SEM from 3 independent experiments. (D) Quantification by BrdU assay of the percentage of cells in each phase of the cell cycle for siRNA-transfected HUVEC or treated with blebbistatin or Y27632. Error bars are means ± SEM from 5 (drug treatments) and 8 (ROCKs silencing) independent experiments. (E) Quantification of the percentage of cells positive for Ki-67 staining. Error bars are means ± SEM from 5 (drug treatments) to 4 (ROCKs silencing) independent experiments. (F) Heatmap of expression of ECM remodelling proteins (left) and of SASP factors (right) over the 4 siRNA conditions, 3 replicates per condition. (*) P-value<0.05; (**) P-value<0.005; (***) P-value<0.0005; (****) P-value<0.00005. Scale bars are equal to 10 µm. Data for siNT and siCCM2 HUVEC are the same as in figure 2.

Our next goal was then to know whether the mechanical defects provoked by dysregulated ROCKs play a direct role in the premature senescence of CCM2-depleted HUVEC. To answer this question, we treated siCCM2 HUVEC with blebbistatin, an inhibitor of myosin II that blocks cell contractility or with Y27632, an inhibitor of ROCK1 and 2 kinase activities. Both treatments inhibited the production of transversal actin stress fibers by siCCM2 HUVEC and restored a more cortical actin rim alike the one observed in control HUVEC (Fig S3A). Interestingly, these treatments inhibited all the senescent traits studied above, i.e. SA-β-gal activity, HIRA-positive SAHF and accumulation in G1 phase of the cell cycle and restored normal level of Ki67+ cells (fig 5A, B, D, E) supporting the fact that increased contractility is involved in the premature senescence of CCM2-depleted EC. Noticeably, blebbistatin and Y27632 had no effect on siNT HUVEC in these assays (Fig S3B, C, and D).

Overall, these data reveal that dysregulated ROCK1 and ROCK2 functions together with increased cell contractility lead CCM2-depleted EC to enter into a premature senescence.

### 5) ROCK1 causes ECM degradation and supports invasiveness of CCM2-depleted EC and neighbouring WT EC

Cancer cells undergoing SASP can promote tumour development through a juxtacrine effect on their microenvironment by secreting matrix metalloproteinases (MMPs) and ECM remodeling enzymes that facilitate tumour cell invasiveness and metastasis(42). Similar to cancer progression, the formation of CCM lesions could result from a SASP-dependent invasion of the brain tissue by EC. Consistently, MMPs have been found around CCM lesions in human(14) and in mouse(13), or zebrafish models(22) and their upregulation plays a role in CCM defects (12,20). To know whether upregulated expression of SASP factors confers invasive skills to siCCM2 HUVEC, we tested the ability of these cells to degrade ECM and invade a 3D matrix. To visualize the degradation of the ECM, siRNA transfected HUVEC were cultured overnight on fluorescent gelatin. siNT HUVEC barely degraded the gelatin as expected for differentiated EC (Fig 6A). Conversely and consistently with their new SASP expression program, siCCM2 HUVEC degraded the gelatin through scratch zones appearing dark in the fluorescently labelled layer (Fig 6A). The linear shapes of these scratch zones suggested that they were produced under focal adhesions. They were dependent on the activity of MMPs as demonstrated by their complete disappearance upon MMP inhibitor GM-6001 treatment (Fig 6A). Together with blocking the mechanosensitive assembly of focal adhesions (Fig S4), Y27632 and blebbistatin blocked the degradation of gelatin (Fig 6A). We previously showed that additional silencing of ROCK1, but not of ROCK2, limits excessive focal adhesion formation in siCCM2 HUVEC(29). Accordingly, only the additional depletion of ROCK1 but not ROCK2 inhibited ECM degradation by siCCM2 HUVEC (Fig 6A). Overall, these data show that ROCK1-dependent increase in cell contractility and in cell-ECM adhesive sites together with the overexpression of MMPs are responsible for the acquisition of ECM degradative skills by siCCM2 HUVEC.

**Figure 6.**
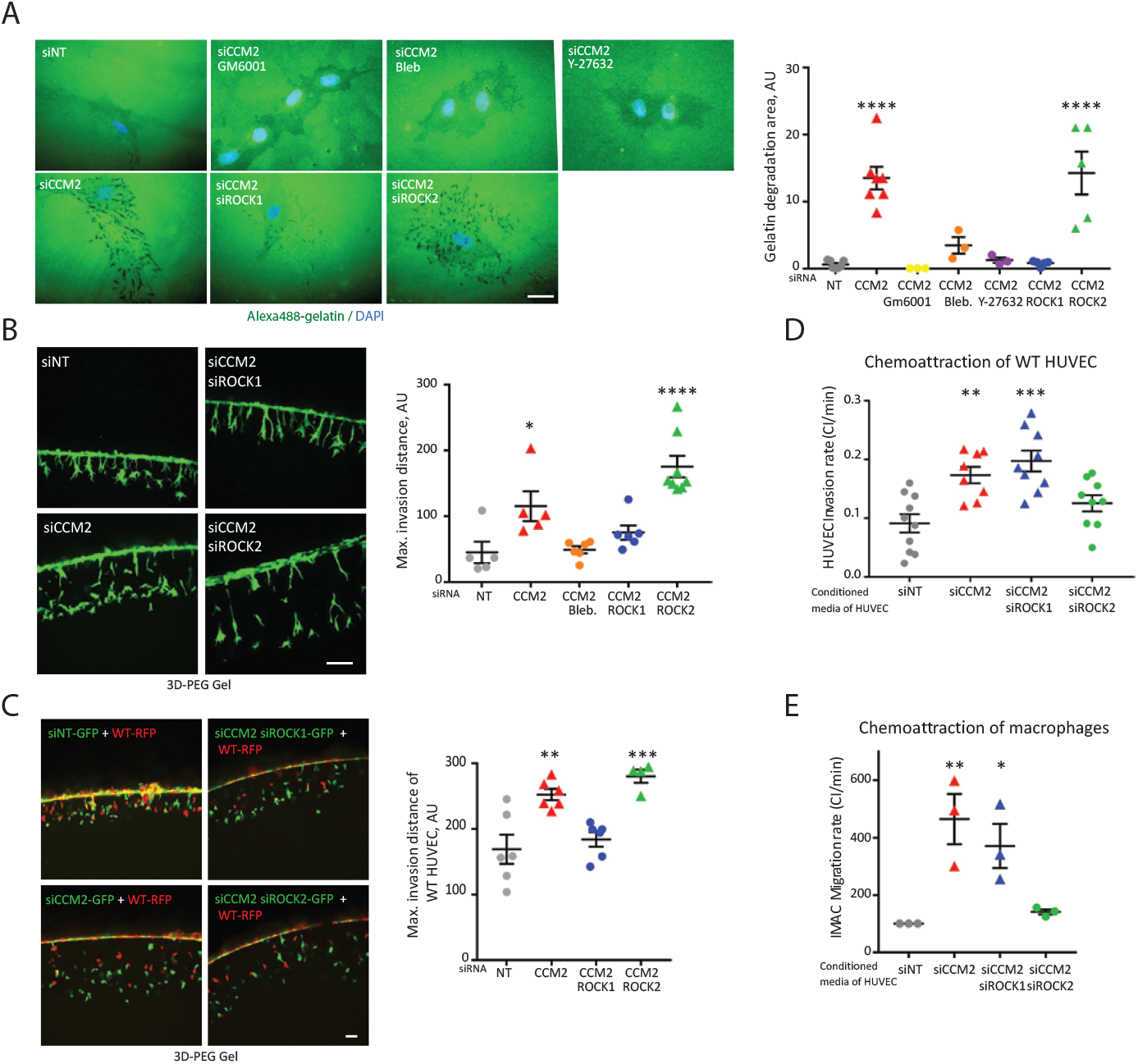
ROCK1 causes ECM degradation and invasion by CCM2-depleted HUVEC and neighbouring WT EC while ROCK2 causes chemo-attraction of WT EC and macrophages. (A)(left) Representative images of the degradation of fluorescent gelatin by siRNA transfected HUVEC or treated with GM6001, blebbistatin or Y27632. Scale bars, 10µm. (right) Quantification of the area of gelatin degradation. Error bars are means ± SEM from 3 (drug treatments) or 5 (silencing of ROCKs) independent experiments. (B)(left) Representative images of siRNA transfected GFP-HUVEC after invasion of 3D-PEG gels. Scale bar, 50µm. (right) Quantification of the maximum invasion distance of siRNA transfected HUVEC or treated with blebbistatin. Error bars are means ± SEM from 3 independent experiments (2 to 3 technical replicates per condition) for siRNA transfected HUVEC and 2 for blebbistatin treated HUVEC. (C)(left) Representative images of siRNA transfected GFP-HUVEC and RFP WT-HUVEC after invasion of 3D-PEG gels Scale bar, 50µm. (right) Quantification of the maximum invasion distance of RFP WT-HUVEC. Error bars are means ± SEM from 3 independent experiments (2 to 3 technical replicates per condition). (D) Quantification of the rate of transmigration of IMAC macrophages measured in a modified Boyden chamber in real time using xCELLigence upon chemo-attraction by conditioned media of siRNA-transfected cells. Error bars are means± SEM from 3 independent experiments. (E) Quantification of the rate of invasion of WT HUVEC measured in a modified Boyden chamber in real time using xCELLigence upon chemo-attraction by conditioned media of siRNA-transfected cells. Error bars are means± SEM from 3 independent experiments (2-4 technical replicates per condition). (*) P-value<0.05; (**) P-value<0.005; (***) P-value<0.0005; (****) P-value<0.00005.

We next tested whether their new degradative capacities gave invasive skills to siCCM2 HUVEC. To measure 3D invasiveness, GFP-expressing siRNA treated HUVEC were plated on 3D-degradable polyethylene glycol gels. Invasive sprouts were imaged after 18 hours and the maximum invasion distance was quantified. siCCM2 HUVEC invaded the 3D gel twice as deep as siNT HUVEC and they mostly invaded as isolated cells with filopodia at their front compared to the cohesive invasion mode of siNT HUVEC (Fig 6B). Silencing of ROCK1 or blebbistatin treatment reduced the invasiveness of siCCM2 HUVEC to the level of siNT HUVEC whereas silencing of ROCK2 enhanced it (Fig 6B) consistently with the increased traction forces upon additional depletion of ROCK2(29).

CCM lesions are mosaics of mutant and WT EC as shown in human and murine lesions(9–11). Moreover, it has recently been shown that CCM3 KO EC can attract WT EC *in vivo* in the brain vasculature and *in vitro* on mixed monolayers(11). Therefore, we asked whether senescent siCCM2 HUVEC could also stimulate the sprouting of WT HUVEC when mixed together on the surface of 3D-PEG gels. Strikingly, we observed that RFP-expressing WT HUVEC invaded the gels twice as deep when they were mixed with GFP-expressing siCCM2 HUVEC as when mixed with GFP-expressing siNT HUVEC (Fig 6C). Interestingly, this increased invasion was significantly reduced when WT HUVEC were mixed with siCCM2+siROCK1 HUVEC but remained unchanged when they were mixed with siCCM2+siROCK2 HUVEC (Fig 6C).

Altogether, our results show that senescent siCCM2 HUVEC have acquired a ROCK1-dependent capacity to invade the ECM and sustain the invasion by WT HUVEC.

### 6) ROCK2 causes the expression of paracrine factors by CCM2-depleted EC that chemo-attract WT EC and immune cells

Apart from remodelling their surrounding ECM, senescent cells secrete paracrine factors that have been shown to promote invasiveness of neighbouring cells and engage deleterious inflammatory responses (42). Therefore, we tested whether secreted factors would have a long distance paracrine effect allowing the recruitment of WT EC. We measured the capacity of a media conditioned by siCCM2 HUVEC to chemo-attract serum-starved WT HUVEC through a layer of Matrigel in a modified Boyden chamber measuring the impedance of cells. Remarkably, siCCM2-conditioned media significantly increased the speed of invasion of WT HUVEC through the Matrigel showing that chemo-attractive factors secreted by siCCM2 HUVEC could also attract EC (Fig 6D). Interestingly, this effect was inhibited by ROCK2 but not ROCK1 additional silencing (Fig 6D). Therefore, siCCM2 HUVEC produce paracrine factors that attract WT EC in a ROCK2-dependent manner.

Since a chronic inflammation is observed at the site of human and murine CCM lesions through the recruitment of activated lymphocytes and monocytes(45), we tested whether the loss of CCM2 leads to the secretion of chemo-attractive cues for immune cells. Thus, we measured the capacity of media conditioned by siCCM2 HUVEC to chemo-attract IMAC, an immortalized macrophage cell line in a modified Boyden chamber as above. Almost no transmigration of macrophages was observed in the case of siNT-conditioned media (Fig 6E). Strikingly, siCCM2-conditioned media provoked a rapid transmigration of the macrophages suggesting that siCCM2 HUVEC had secreted chemo-attractive factors that siNT HUVEC had not (Fig 6E). Importantly, this chemo-attraction was inhibited when ROCK2 but not ROCK1 was additionally silenced (Fig 6E). Therefore, siCCM2 HUVEC secrete paracrine factors that can attract immune cells such as macrophages. Moreover, these chemo-attraction skills depend on ROCK2.

Overall, these results demonstrate for the first time to our knowledge that, when CCM2 is lost, HUVEC undergo a SASP that is driven by the mechanical and molecular disorders provoked by ROCKs dysfunctions. A major consequence of this SASP is a profound mechano-dependent remodelling of the microenvironment leading to the recruitment of wild-type EC and immune cells to generate mosaic CCM lesions.

## Discussion

Many cellular pathways are dysregulated in CCM-deficient EC and it has been difficult to propose a mechanistic model that could take into account all these aspects. Transition in cellular fate(18) upon morphological and mechanical changes during the loss of cell-cell junctions(19,29,46) are associated with overproduction of ROS(16), decreased autophagy(17), secretion of metalloproteases(12–14) or cytokines(15) and increased integrin(19,20), p38(21), MEKK3/KLF2(20,22,23) and TLR4(24) signaling. The first breakthrough of this current study is to show, for the first time to our knowledge that CCM2-deficient EC engage towards a Senescence Associated Secretory Phenotype (SASP) (Fig 7). Multiple senescent traits were validated such as transcriptomic SASP signature, lysosomal SA-β-galactosidase activity, upregulation of CDK inhibitors, cell cycle blockage in G1, presence of SAHF along with the secretion of chemo-attractant factors. Moreover, we show that the functional consequence of this SASP is an acquired ability of CCM2-deficient EC to invade the ECM and recruit wild-type EC and immune cells.

**Figure 7.**
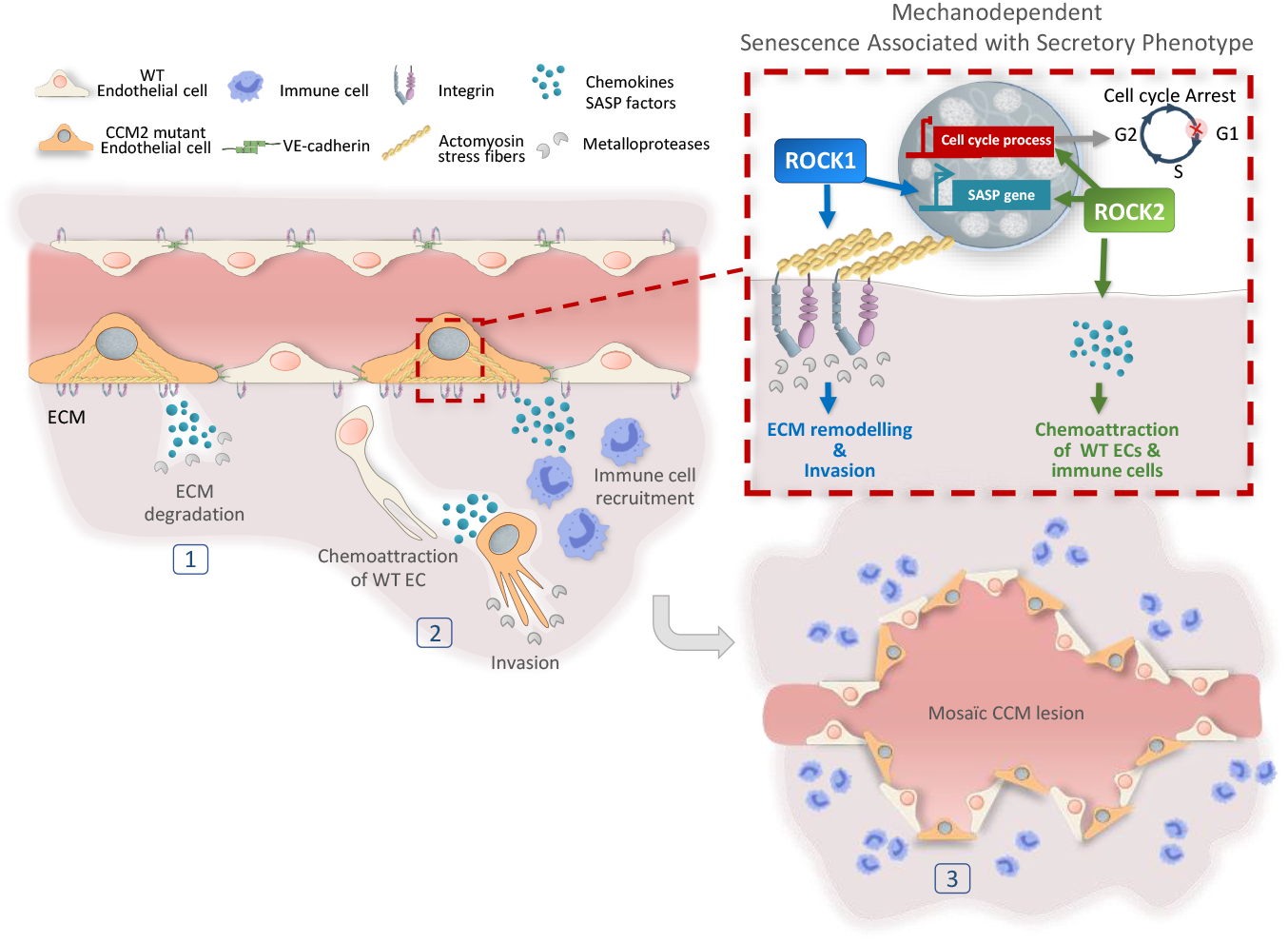
Proposed mechanism for CCM2 lesion mosaicism: a mechano-dependent SASP with complementary roles for ROCK1 and ROCK2. In low shear stress capillaries, the loss of CCM2 protein upon the second-hit mutation induces a ROCK-dependent premature senescence of the mutant endothelial cell associated with secretory phenotype (SASP). (1) This senescent cell acquires ECM degradative skills by secreting metalloproteases. (2) It invades the surrounding tissue, providing tracks for neighbouring WT EC and produces SASP factors that chemo-attract WT EC and immune cells. (3) These processes allow the formation and expansion of a mosaic CCM lesion. Dysregulated ROCKs have complementary key roles. Whereas, they both control the expression of genes involved in the SASP, with ROCK2 having a more prominent regulatory role, ROCK1 is specifically required for ECM invasion by mutant and WT EC. ROCK2 is specifically required for paracrine chemo-attraction of WT EC and immune cells.

SASP is characterized by cell growth arrest, widespread changes in chromatin organization and gene expression(36). These changes also include the secretion of numerous pro-inflammatory cytokines, chemokines, growth factors and proteases(43). Acute senescence is beneficial in development, tissue regeneration or cancer through the clearance of senescent cells by the immune system(47). Chronic senescence triggers chronic inflammation that can on the contrary favour age-related diseases including cancer, fibrosis, cardiovascular diseases, type 2 diabetes, osteoarthritis or neurological disorders(33). Indeed, the long-term secreted pro-inflammatory factors promote cell proliferation, microenvironment remodelling, angiogenesis and inflammation in a paracrine manner. Remarkably, decreased autophagy(48), ROS overproduction(49), P38 MAPK(50), KLF2/4(51,52) and TLR4(53) signaling pathways have all been involved in the induction of SASP either through regulation of CDK inhibitors or through the NFkB or C/EBPβ-driven expression of cytokines. Therefore, we propose that each of the dysregulated features of CCM-deficient EC represents a different facet of the same cellular state. Future research should help reconstructing the complex chronology of the different events in the framework of this SASP. This should identify key therapeutic targets for either single or combinatorial drug treatments to block at their root the defective molecular pathways involved in the CCM2 disease. Senolytic drugs and drugs targeting the SASP *per se* are currently under pre-clinicial trials for cancer therapy or other age-related diseases(33). They could be tested to prevent the formation and expansion of CCM lesions. A recent study strongly supports that premature aging of the neuro-vascular system could be the cause of CCM disease by showing that aging and CCM brains share common dysregulated features including impaired endothelial barrier function, inflammation, 320 DEGs, plasma molecules and imaging biomarkers (54).

This SASP not only reconciles all the CCM defects but also brings a molecular explanation for the mosaicism of the CCM lesion (Fig 7). Secreted chemokines and cytokines attract wild-type EC in paracrine and juxtacrine manners and trigger an inflammatory response, another major feature of CCM lesions(11). Not only are WT EC attracted by CCM-deficient EC but they are triggered to undergo EndMT and express stem cell markers(11). Interestingly, NFkB signalling has been shown to activate WNT/βcatenin signalling to induce the EMT of cells into cancer stem cells(55) and to induce cancer cell metastasis in response to IL1β(56). Therefore, one can assume that a similar mechanism might operate in CCM lesion where senescent cells could induce the dedifferentiation of WT EC through EndMT.

Whereas it has also been observed that Ccm3-null EC attract wild-type EC at the site of the lesion in Ccm3 mouse model, clonal expansion of Ccm3-null EC precedes the expansion of the lesion(10,11). Moreover, differently from KRIT and CCM2, the loss of CCM3 actually preserves primary EC from replicative senescence *in vitro*(57) which is consistent with our findings. Therefore, opposite mechanisms may lead to the formation of CCM lesions upon the loss of CCM3 and CCM2 or KRIT. A rapid clonal proliferation of CCM3-deficient EC may trigger the formation of aggressive lesions at early age whereas an entry in SASP of KRIT- or CCM2-deficient EC may lead to the progressive maturation of lesions and to the later onset of the CCM1/2 disease. Accordingly, the absence of increased Ki67 or phospho-histone 3 staining in the EC lining the lesions in the acute Ccm2^-/-^(59) and in the Ccm1^+/-^; Msh2^-/-^ (7) mouse models or in ccm1^-/-^mutant zebrafish embryo(60) do not argue in favour of an early clonal endothelial proliferation upon CCM1 or CCM2 loss. Studies are ongoing in several laboratories to better characterize the different cellular states that co-exist in mutant and WT EC populations over time within the CCM lesions (58). However, the complexity of the situation renders these *in vivo* investigations very challenging and do not tackle the relationship between mechanotransduction and CCM progression. This current study on pure or mixed populations of CCM2-depleted and WT EC has led to significant insights towards a better understanding of the complex *in vivo* situation. It should constitute the foundation for new specific *in vivo* studies. Using the confetti reporter system in Ccm1/2 mouse models compatible with SA-βgalactosidase staining would help studying the initiation and expansion of the lesions in comparison with the Ccm3 model.

The second major breakthrough of this study is that CCM proteins are involved in a mechanotransduction pathway controlling the onset of the SASP. The fact that cellular mechanical defects can provoke senescence is an emerging area of investigations. Not only does our study make a clear demonstration of it, but it also identifies CCM proteins and ROCKs as part of a crucial scaffold controlling senescence. We previously showed that upon the loss of CCM1/2, a vicious cycle sets up between aberrant ECM remodelling and increased intracellular contractility which breaks the permeability barrier of the endothelium(19). Moreover, we demonstrated that placing WT EC on ECM remodelled by mutant EC provoke CCM-like morphological defects in these cells(19). Going deeper into mechanistic investigations, we showed that upon the loss of CCM2, ROCK2 is delocalized from VE-cadherin junctions while ROCK1 gets over activated, enhancing the polymerization of β1 integrin-anchored stress fibers(29). Here we uncover for the first time that, beyond their role in the disruption of the endothelial architecture, dysregulated ROCKs are instrumental in the transition of CCM2-depleted EC towards SASP (Fig 7). We propose that this SASP allows the amplification of local mechanical perturbations into a systemic response that propagates to other EC and immune cells, a phenomenon that could also exist for other mechano-dependent pathologies such as cancer or fibrosis.

ROCK2 controls the expression of a very significant fraction of the genes dysregulated upon the loss of CCM2 among which genes involved in senescence, cell cycle and paracrine SASP factors. The opposite expression profiles in absence of CCM2 or ROCK2 suggest that ROCK2 gains a broad transcriptional regulatory function when its scaffold CCM2 is lost. In fact, ROCK2 has already been involved in cellular senescence(61). In this study, the loss of the two ROCK isoforms led to senescence of MEF and was due to the downregulation of CDK1, CKS1 and cyclin2A. Intriguingly, we observe a similar downregulation of these genes in HUVEC when CCM2 is lost (Fig S2B), but contrary to this other study, their expression is rescued by the additional silencing of ROCK2. This suggests that the presence or absence of CCM2 has a strong influence on the transcriptional activity of ROCK2. Future mechanistic studies will be necessary to understand how ROCKs interplay with the CCM complex to regulate gene expression.

We previously showed that ROCK1 over activation allows CCM2-depleted HUVEC to exert stronger traction forces on the ECM (19,29). Here we show that ROCK1 enables the mechanosensitive degradation and invasion of the ECM by the senescent CCM2-depleted EC and by neighbouring WT EC. Since EC pull on their microenvironment to invade it(62), it is likely that these stronger forces are responsible for enhanced invasion. Further investigations will help better understand the molecular events at play. Like ROCK2, ROCK1 affects gene expression downstream of CCM2 though to a lesser extent. Moreover, inhibition of the myosin motor or silencing of ROCK1 is enough to block senescence. These data suggest that cross talks exist between the contractility of the acto-myosin cytoskeleton and the transcription of genes involved in SASP. Several mechanosensitive transcription factors are known to be regulated by acto-myosin polymerization or contractility(63). Moreover, ROCK2 can shuttle into the nucleus where it phosphorylates the chromatin remodeller p300HAT(64). Future investigations should unravel how ROCK1 and ROCK2 cooperate to regulate chromatin organization and gene expression through the control of mechanosensitive transcription machineries.

Overall, this study demonstrates that CCM2-deficient EC undergo a SASP that is driven by the mechanical and molecular disorders provoked by ROCKs dysfunctions. Remarkably, this discovery unifies all the known dysregulated traits of CCM1/2-deficient EC into a unique cellular state. A major consequence of this mechano-dependent SASP is the remodelling of a microenvironment that sustains chronic recruitment of wild-type EC and immune cells to the site of the lesion. Thus, this work unravels the molecular basis of the mosaicism of CCM1/2 lesions and their inflammatory status. It opens new avenues for mechanistic *in vivo* studies on the dynamics and penetrance of the CCM1/2 disease as well as for exploring new therapies.

## Material and Methods

### Cell culture and transfections

Pooled HUVEC were obtained from Lonza. Upon reception, HUVEC at P0 were expanded over 2 passages in collagen 1 (from rat tail, BD) coated flasks in complete EGM-2 medium supplemented with 100 U/ml penicillin and 100 μg/ml streptomycin at 37°C in a 5% CO2 - 3% O2 humidified chamber. HUVEC at passage 3 were transfected twice at 24 h-intervals with 30 nM siRNA and Lipofectamine RNAi max (Life Technologies, ref. 13778-150) according to the manufacturer’s instructions in 37°C in a regular 5% CO2 in a humidified chamber. For double transfections, 30 nM of each siRNA duplexes (Dharmacon smartpool ON-TARGET plus Thermo Scientific) was used; Non-targeting siRNA #1, CCM2; ref. L-014728-01), KRIT; ref. L-003825-00, CCM3; ref. L-004436-00, ROCK1; ref. L-003536-00 and ROCK2; ref. L-004610-00. When required, blebbistatin, Y27632 and GM6001 were used at 10 μM final.

### RNA sequencing and differential analysis

One million of siRNA transfected HUVEC were seeded at confluency the day after the second round of transfection in wells of 6-well plates coated with collagen 1 (from rat tail) and cultured for 48h in complete media at 37°C, 5% CO2. Total RNA were extracted from HUVEC using the NucleoSpin RNA II kit (Macherey-Nagel) according to the manufacturer’s instructions. cDNA libraries were prepared with the TruSeq Stranded mRNA Sample Preparation (Illumina) and sequenced on a HiSeq 2500 Illumina platform using single-end 50 basepair reads at the MGX facility (Montpellier). Fastq files were aligned using STAR (2.5.2b) on UCSC hg19 genome. The contents of the annotation directories were downloaded from UCSC on: July 17, 2015. Bam files were counted using htseq-count (0.11.2.) with option -t exon -f bam -r pos -s reverse -m intersection-strict -nonunique none. Differential analyses were performed with SARTools (1.4.1) (65) using DESeq2 (1.12.3) (66) and default options. P-values were adjusted with Benjamini-Hochberg procedure(35) set for 5% of FDR. Heatmaps using a correlation matrix and boxplots were obtained with custom R script (3.3.0). The raw sequencing data used in this study are available in the National Center for Biotechnology Information’s Gene Expression Omnibus (GEO) database and are accessible through GEO series accession number (pending).

### Data Availability

The raw data (FastQ files) and processed data (count files) are deposited in the Gene Expression Omnibus database under ID code GEO: GSE165406.

### Bioinformatics analyses

Gene ontology analyses on DEG in siCCM2 condition (fold change [FC] ≥ 2; *P* < 0.05 with FDR corrected using Benjamini Horchberg method (35)) were made with PANTHER version 15,0 Released 2020-02-14 using slim cellular components and biological processes. Enriched pathways analyses were conducted with Reactome. DEGs (*P* < 0.05, 5% FDR) rescued by additional silencing of ROCK1 or ROCK2 were recovered at the union of Venn diagrams of DEGs in siCCM2 vs siNT and DEGs in siCCM2 vs siCCM2+ROCK1 or ROCK2 respectively. Gene Set Enrichment Analyses were conducted using GSEA software from Broad Institute, UC San Diego(67).

### Quantitative RT-PCR

Purified RNA (1 μg) were reverse transcribed using the iScript Reverse Transcription Supermix (Biorad). Quantitative real-time PCR (Q-PCR) was performed with iTaq Universal SYBR Green Supermix (Biorad) in a 20 μl reaction on a thermal cycler (C-1000 Touch; Bio-Rad Laboratories). Product sizes were controlled by DNA gel electrophoresis and the melt curves were evaluated using CFX Manager (Bio-Rad Laboratories). A total of three housekeeping genes were selected for their stability in our HUVEC cell line under our experimental conditions, using the three analytical software programs, geNorm, Normfinder and Bestkeeper (68,69). We used the relative expression software tool CFX Manager for relative quantification, and normalization was achieved using a normalization factor from all reference genes(68). The mean of three technical replicates was calculated per biological replicate.

### Immunofluorescence staining

HUVEC were seeded at 5×10^4^ cells or 2×10^5^ cells in 24-well plates on coverslips coated with 10 μg/ml fibronectin (from human plasma, Sigma Aldrich) and incubated overnight in complete EBM-2 medium. Cells were fixed with 4% PFA, permeabilized with 0.2% Triton X-100, and incubated with anti-activated β1 integrin clone 9EG7 ((BD Biosciences, 1/200), anti-Ki67 (AN9260 Millipore, 1/200), anti-HIRA (WC119, Millipore, 1/200) antibodies. After rinsing, coverslips were incubated in Goat anti-Mouse or anti-Rabbit IgG (H+L) highly cross-adsorbed secondary antibody, Alexa Fluor conjugated AF 488, AF 546, AF 647 (Invitrogen, 1/1000) and phalloidin conjugates with Atto 647 (Sigma, 1/2000). The coverslips were mounted in Mowiol/DAPI solution and imaged on an epifluorescent Axiomager microscope (Zeiss) at 63X magnification.

### SA-B-Galactosidase staining

HUVEC were seeded 48 hours after the second siRNA transfection at a density of 5×10^4^ in 24-well plates coated with 10µg/mL fibronectin and incubated overnight in complete EBM-2-medium. Senescence-Associated β-galactosidase activity was assessed using a SA-β-galactosidase staining kit according to the manufacturer’s instructions (Cell Signaling). Positive cells were counted manually out of more than 100 cells total.

### BrdU assay

HUVEC were seeded 48 hours after the second siRNA transfection at a density of 2×10^5^ cells in 6-well plates coated with 100µg/mL collagen 1 and incubated overnight in complete EBM-2 medium. The BrdU assay was performed with the BD Accuri C6 flow cytometer using the APC-BrdU flow kit according to the manufacturer’s instructions (Cell Signaling).

### xCELLigence proliferation assay

Proliferation assay was performed using the xCELLigence Real-Time Cell Analysis (RTCA) DP instrument in combination with E-plate 16 (ACEA Biosciences) coated with 100µL of 100µg/mL collagen 1 (from rat tail) for 30 min at 37°C and washed 2 times with PBS 1X. 40 µL of complete EBM-2 medium was added and baseline without cells was made with RTCA software. 5×10^3^ siRNA transfected HUVEC were seeded 48h after the second round of transfection in 100µL of complete EBM-2-medium. Impedance measurements were recorded every 5 min during 24h. Impedance was normalized at 4 hours after seeding to eliminate the contribution of cell spreading and adhesion to the signal. Slope measurement was performed between 4h and 24h.

### Matrigel invasion assay

This assay was performed using the xCELLigence RTCA DP instrument in combination with CIM-Plate 16 (ACEA Biosciences). A layer of 3.3% of growth factor reduced Matrigel (Corning) diluted with EBM basal media was poured on ice in the upper chamber and incubated overnight at 37°C for polymerization. 40 μl basal EBM media was added to the upper chamber and the baseline made after equilibration at 37°C. After setting the baseline, 3×10^4^ WT HUVEC were added to the upper chamber. The conditioned media from siRNA transfected cells was recovered 48 h after the second round of transfection and centrifuged at 12000 rpm for 5 min and 160µL was added to the lower chamber of the CIM-plate. After one hour of equilibration at 37°C, impedance was measured every 3 minutes for 24 hours.

### Chemo-attraction assay

This assay was performed using the xCELLigence Real-Time Cell Analysis (RTCA) DP instrument in combination with CIM-Plate 16 (ACEA Biosciences). After setting the baseline, 3×10^4^ IMAC were added in 100µL of basal EBM in the upper chamber and 48h-conditioned media from siRNA transfected cells added to the lower chamber of the CIM-plate. After 30 min equilibration at 37°C, impedance was measured every 3 minutes for 24 hours.

### Gelation degradation assay

Coverslips in 24 well-plates were coated with gelatin-Alexa488 dye as previously described (70). SiRNA transfected HUVEC were seeded 48h after the second siRNA transfection at 5×10^4^/well. HUVEC were incubated in OptiMEM medium overnight at 37°C in a 5% CO2 incubator, fixed with 4% PFA, and washed with PBS thrice. For a quantitative analysis of the degradative skill of siRNA transfected cells, 5 images per condition were randomly acquired with an epifluorescent Axiomager microscope (Zeiss) at 40X magnification, and were convertedto binary images using B&W thresholding on ImageJ. The total area of the black zones corresponding to the total area of degradation of the fluorescent gelatin was measured.

### 3D-PEG invasion assays

Poly-ethylene glycol (PEG) hydrogels were prepared on ice in EBM-2 complete media by combining an MMP-sensitive peptide modified PEG precursor (8-arm 40kDa,(71)) at 1.5% polymer concentration, 50 μM Lys-RGD peptide (Pepmic), and 1 μM sphingosine-1-phosphate (S1P, Sigma-Aldrich). The hydrogels were enzymatically crosslinked using a reconstituted and thrombin-activated Factor XIII (Fibrogammin, CSL Behring), prepared as previously described (72)) at 10% of total hydrogel volume. A 20 μL volume of the hydrogel suspension was pipetted into a modified imaging chamber (Secure-Seal™ hybridization sealing systems, ThermoFisher Scientific) attached to the bottom of a 24-well plate held vertically (62). The hydrogel was allowed to polymerize for 30 minutes in this orientation at room temperature prior to cell seeding. Depending on the study, a confluent cell monolayer composed of either 5×10^4^ siRNA transfected GFP-HUVEC or a 1:1 ratio mixture of siRNA transfected GFP-HUVEC and naїve RFP-HUVEC was allowed to adhere to the PEG meniscus at 37°C, 5% CO2 for 1 hour and then placed back horizontally with 1mL of EBM-2 complete media. Fixation with 4% PFA in DPBS was performed after 24 hours. Sprouts invading the PEG hydrogel were imaged using a Leica SP8 inverted confocal microscope with an HC PL APO 10x, 0.4 numerical aperture dry objective to obtain image stacks at 1024×1024 pixels with a 50 μm Z-stack at 1-1.5 μm Z-spacing. A Z-projection of each image was used to manually quantify invasion distances using the line measurement tool of ImageJ. More than 100 sprouts were analyzed per technical replicate.

### Statistical tests

Results were assessed by either performing a paired t-test for comparing 2 conditions or for more than 2 conditions by the Tukey’s multiple comparison tests post-ANOVA to compare with control; a 0.5 alpha level was used for all comparisons. Prism software was used to conduct the statistical analysis of all data. P<0.05 was considered to be significant. *P<0.05, **P<0.05, ***P<0.005. n represents biological replicates.

## Acknowledgements

Most of the computations presented in this paper were performed using the CIMENT/GRICAD infrastructure (https://gricad.univ-grenoble-alpes.fr). The authors acknowledge the EpiMed core facility (http://epimed.univ-grenoble-alpes.fr) for their support and assistance in this work. This study was supported by the ANR (ANR-17-CE13-022), the Fondation pour la Recherche Médicale FRM (DEQ20170336702), the International Emerging Action CNRS, the association Espoir contre le Cancer Isère, the FWO project G087018N, infrastructure grant I009718N, Hercules Foundation (G0H6316N), the European Research Council under the European Union’s Seventh Framework Program (FP7/2007–2013)/ ERC Grant Agreement No. 308223). PhD fellowship grants were supported by ANR and FRM to D.V. and FWO (1S68818N) to A.S. We would like to thank Claudia Röedel, Béatrice Eymin, and Gwénola Boulday for sharing ideas and reagents and Salim Seyfried for critical reading of the manuscript.

## Author contributions

D.R.V., A.S., H.V.O., E.F. conceived the project and designed experiments; D.R.V., A.S., S.M., E.P., E.F. performed experiments; F.C., D.R.V., E.F. analyzed the bioinformatical data; O.D. contributed to scientific discussion; P.R. provided reagents; C.A.R., E.F. and H.V.O. provided fundings; E.F. wrote the manuscript which has been revised by all authors.

## Figure legends

**Table S1: Level of expression of the dysregulated genes in the 6 different conditions of siRNA** (A) DEG with FC≥2 (*P* < 0.05, FDR corrected using Benjamini Horchberg method) in siCCM2 HUVEC. Rescued DEG with FC≥2 upon additional silencing of ROCK1 (B) or ROCK2 (C).

**Table S2: Gene sets of Senescence and SASP used for GSEA analyses**.

---

**Fig S1:**
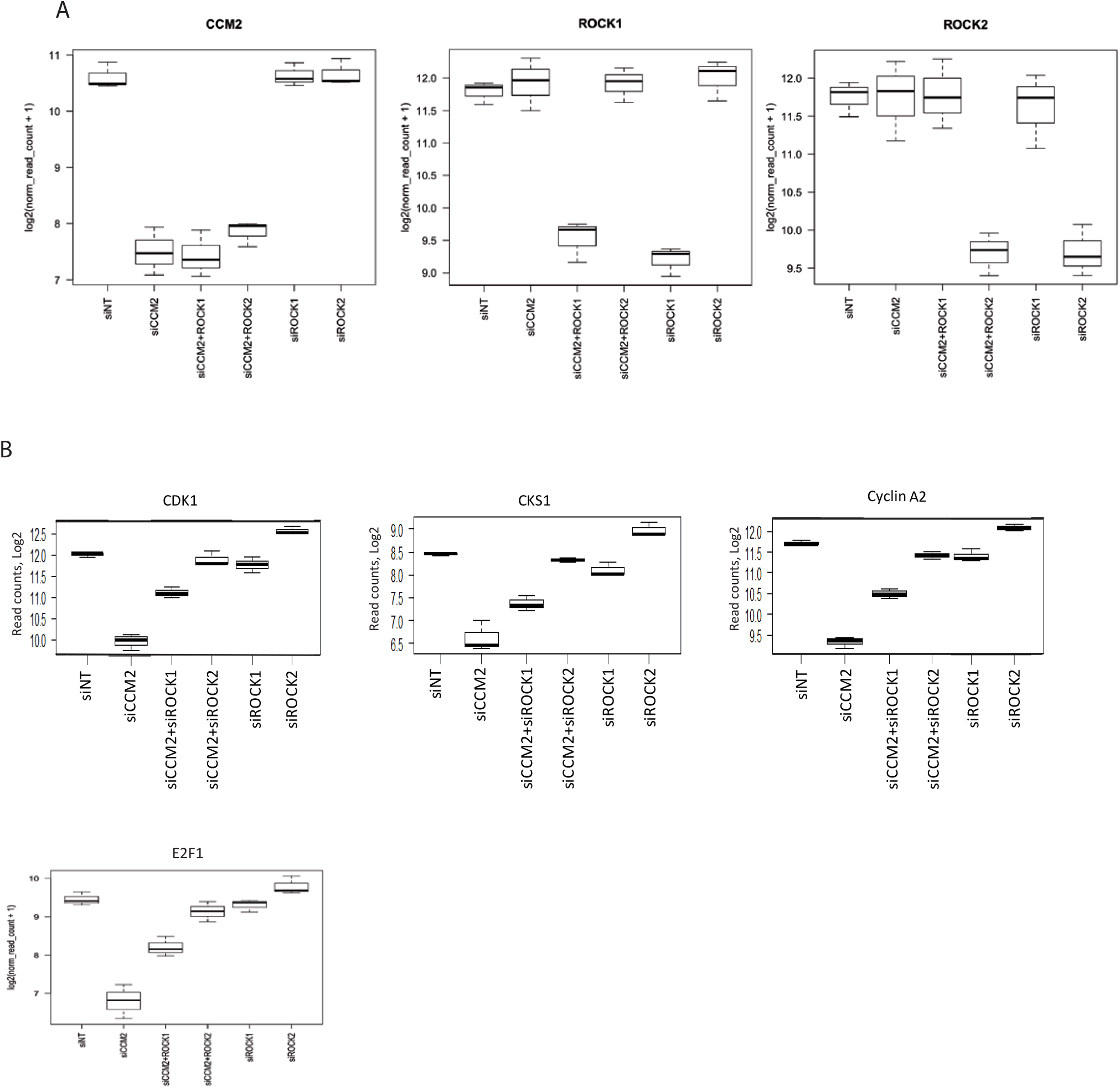
Boxplots of the expression level of CCM2, ROCK1, ROCK2 (A) and cell cycle regulators (B) in the 6 siRNA conditions as measured by RNA seq.

**Fig S2:**
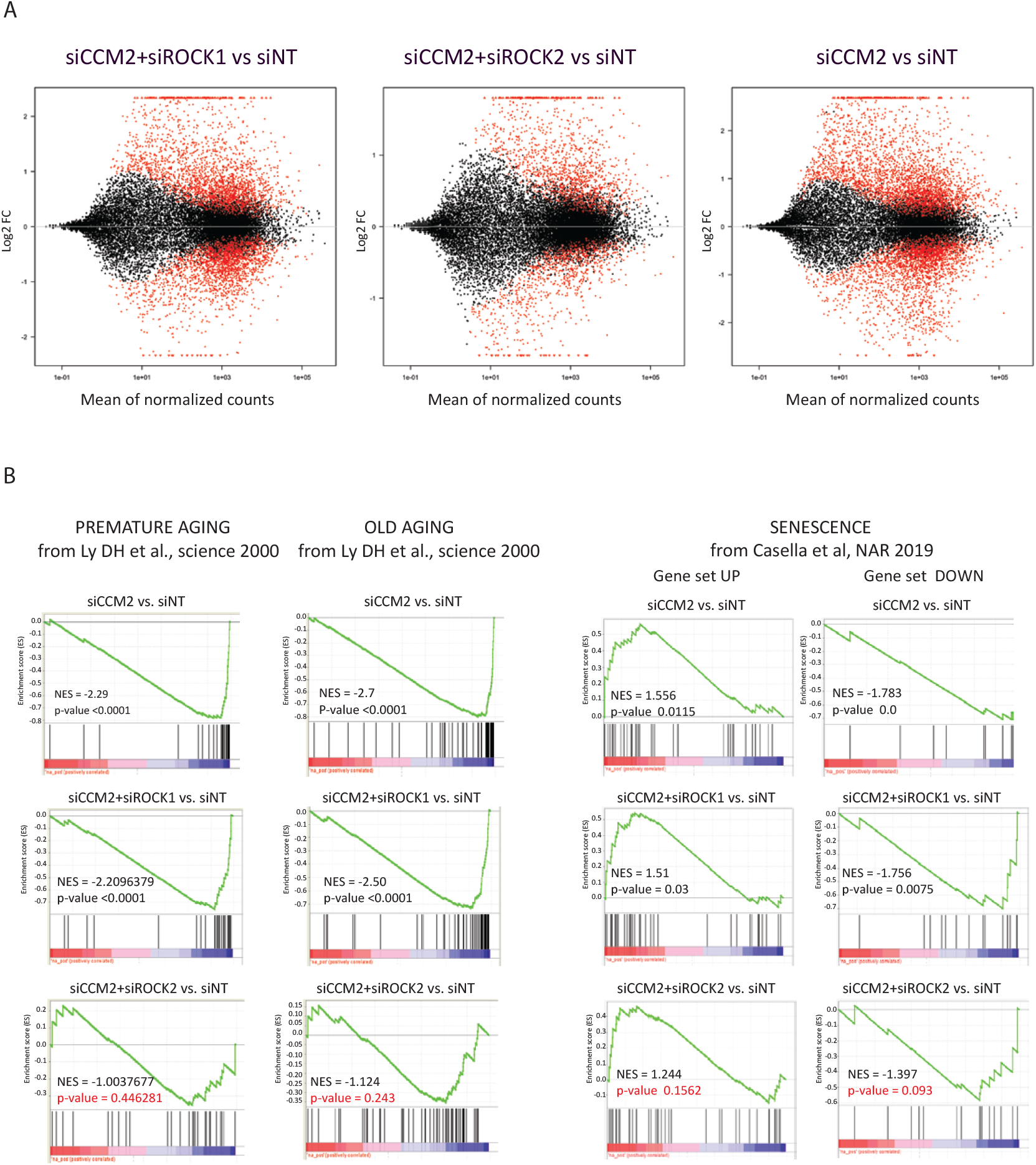
GSEA analyses using different gene sets of senescence. (A) MA-plots showing the the better restoring effect of ROCK2 over ROCK1 on the DEG of CCM2-depleted HUVEC. (B) Comparison of the enrichment in different senescence and SASP signatures(38,40) in siCCM2, siCCM2+ROCK1 and siCCM2+ROCK2.

**Fig S3:**
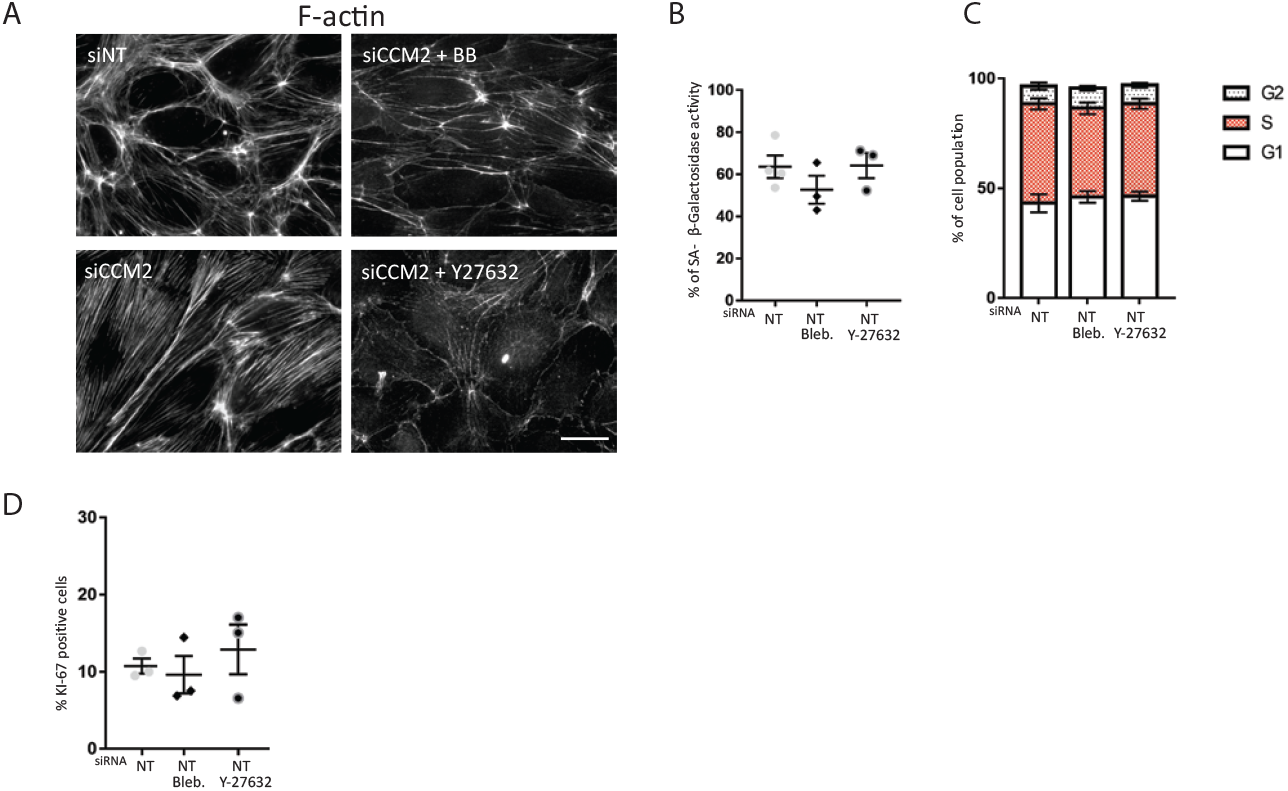
Effect of drugs on the actin cytoskeleton of siCCM2-HUVEC and on marks of senescence of siNT HUVEC. (A) Representative immunofluorescence imagesof the actin cytoskeleton of CCM2 transfected HUVEC or HUVEC treated with drugs, scale bar 10 μm. Effects of drug treatments on SA-β-galactosidase activity (B) cell cycle progression (C) and Ki67 staining (D) of siNT HUVEC.

**Fig S4:**
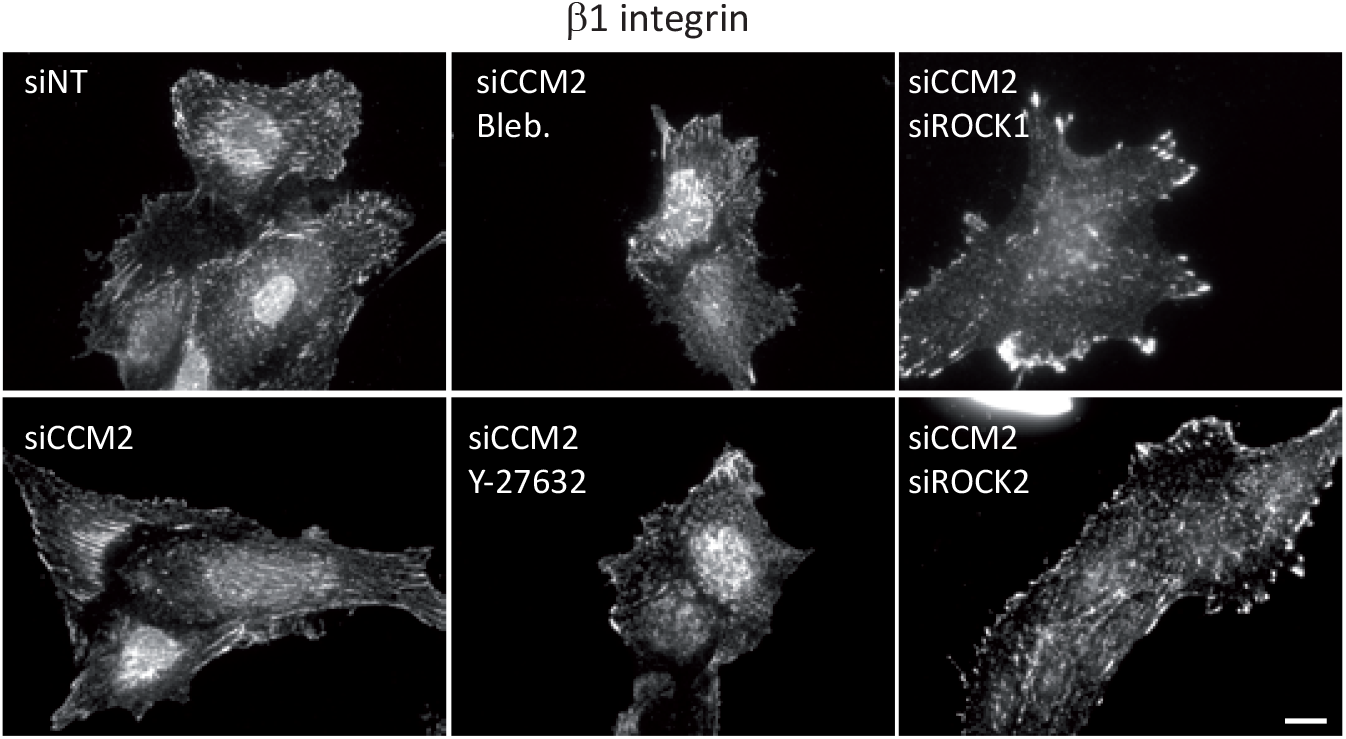
β1 integrin staining of siRNA transfected or drug treated HUVEC spread on gelatin. Scale bar, 10 μm.

